# Mass spectrometry imaging of natural carbonyl products directly from agar-based microbial interactions using 4-APEBA derivatization

**DOI:** 10.1101/2023.05.15.540794

**Authors:** Dušan Veličković, Kevin J. Zemaitis, Arunima Bhattacharjee, Christopher R. Anderton

**Affiliations:** Environmental Molecular Sciences Laboratory, Pacific Northwest National Laboratory, Richland, WA 99354

**Keywords:** MALDI, metabolomics, carboxylic acids, aldehydes, ketones, *Bacillus subtilis*, *Fusarium*

## Abstract

Aliphatic carboxylic acids, aldehydes, and ketones play diverse roles in microbial adaptation to their microenvironment, from excretion as toxins to adaptive metabolites for membrane fluidity. However, the spatial distribution of these molecules throughout biofilms, and how microbes in these environments exchange these molecules remains elusive for many of these bioactive species due to inefficient molecular imaging strategies. Herein, we apply on-tissue chemical derivatization (OTCD) using 4-(2-((4-bromophenethyl)dimethylammonio)ethoxy)benzenaminium bromide (4-APEBA) on a co-culture of a soil bacterium (*Bacillus subtilis* NCIB 3610) and fungus (*Fusarium* sp. DS 682) grown on agar as our model system. Using matrix-assisted laser desorption/ionization mass spectrometry imaging (MALDI-MSI), we spatially resolved more than 300 different metabolites containing carbonyl-groups within this model system. Various spatial patterns are observable of these species, which indicate possible extracellular or intercellular processes of the metabolites, and their up or down regulation during microbial interaction. The unique chemistry of our approach allowed us to bring additional confidence in accurate carbonyl identification, especially when multiple isomeric candidates were possible, and this provided the ability to generate hypotheses about the potential role of some aliphatic carbonyls in this *B. subtilis*/*Fusarium* sp. interaction. The results shown here demonstrate the utility of 4-ABEBA-based OCTD MALDI-MSI in probing interkingdom interactions directly from microbial co-cultures, and these methods will enable future microbial interactions studies with expanded metabolic coverage.

**IMPORTANCE:** The metabolic profiles within microbial biofilms and interkingdom interactions are extremely complex and serve a variety of functions, which include promoting colonization, growth, and survival within competitive and symbiotic environments. However, measuring and differentiating many of these molecules, especially in an *in-situ* fashion, remains a significant analytical challenge. We demonstrate a chemical derivatization strategy that enabled highly sensitive, multiplexed mass spectrometry imaging of over 300 metabolites from a model microbial co-culture. Notably, this approach afforded us to visualize over two dozen classes of ketone-, aldehyde-, and carboxyl-containing molecules, which were previously undetectable from colonies grown on agar. We also demonstrate that this chemical derivatization strategy can enable discrimination of isobaric and isomeric metabolites, without the need for orthogonal separation (*e.g.,* online chromatography or ion mobility). We anticipate this approach will further enhance our knowledge of metabolic regulation within microbiomes and microbial systems used in bioengineering applications.

## INTRODUCTION

Mass spectrometry imaging (MSI) is becoming an established technique for exploring the nature and diversity of the chemical compounds produced in microbial systems (1–3). It has been extensively applied in understanding biochemical processes in microbial and host-microbe interactions (2), biofilm formation (4), and microbial phenotyping (5). Within literature, spatial probing via MSI for retrieving localized metabolic signatures is commonly performed on microbial systems grown on agar or other solid growing medium, including other organisms (6). While there are numerous types of ionization modalities and mass spectrometry approaches such as nanospray desorption electrospray ionization (nanoDESI) (7), laser ablation electrospray ionization (LAESI) (8), liquid extraction surface analysis (LESA) (9), and ultrahigh lateral resolution secondary ion mass spectrometry (SIMS) (10) that have been used to chemically image microbial samples, matrix-assisted laser desorption/ionization (MALDI) methods are most broadly implemented, in part, due to their robustness, reproducibility, high spatial resolution, and wide molecular coverage (1, 11–14).

While there are many classes of metabolites and small molecules that can be readily detected and annotated by MALDI-MSI (14), the ability to obtain comprehensive detection of natural acidic compounds, including aliphatic carboxylates and carbonyls, which play diverse roles in microbes (15–18), and are valuable additives in food, fragrances, and pharmaceuticals (15, 16), remains a significant challenge. For example, MALDI-MSI analyses of agar-based samples has been limited to the positive ionization modality where these classes of biomolecules ionize poorly. The sparse reporting of negative ionization mode analyses is presumed to be ascribed to the chemistry of the agar, which has a negative charge due to sulfate groups. This could dissipate charge during negative ion mode analysis, and/or possibly limit the ability of MALDI matrices used in negative ionization mode to efficiently extract and co-crystalize with analytes on the agar surface. Perhaps this is, in part, a reason why others developed imprinting strategies to transfer colonies from agar to more suitable supports for negative ionization mode analysis (19). Nevertheless, even imprinting approaches have limitations, where transfer efficiency, molecular selectivity, and molecular relocation are notable issues for comprehensive molecular mapping (20).

Here, we provide a new MSI approach for mapping endogenous metabolites containing carbonyl groups from microbial systems. We applied our previously developed on-tissue chemical derivatization (OTCD) protocol to microbial cultures grown on agar as a proof-of-concept of this approach (21). There are a growing number of OTCD reagents and protocols that have been developed to increase the sensitivity and molecular coverage from mammalian and plant samples in MALDI-MSI (22–24). Our approach uses 4-(2-((4-bromophenethyl)dimethylammonium)ethoxy)benzenaminium dibromide (4-APEBA), which adds a permanent positive charge to carbonyl analytes, making them amenable to positive ionization mode analysis MSI, which is especially useful for analyzing agar-based microbial colonies. Additionally, the bromine in 4-APEBA introduces a unique isotopic pattern to derivatized molecules, which can be exploited for more confident analyte annotation (21). We used this approach to study the interaction of soil microbes *B. subtilis* NCIB 3610 and *Fusarium* sp. DS 682 (25). Our results demonstrate that this approach enabled high-sensitivity analysis of over 300 microbially generated carbonyl-containing molecules directly from microbial cultures on agar plates. To our knowledge, this is the first demonstration of using an OTCD approach for MSI-based metabolic profiling of microbial systems.

## RESULTS AND DISCUSSION

Our results show that 4-APEBA-based OTCD of the *B. subtilis* NCIB 3610 and *Fusarium* sp. DS 682 interaction enabled confident chemical formula annotations of over 300 various carbonyls, which putative structural annotations are proposed in **Supplementary Table S1,** and are further classified in **Supplementary Figure S1**. In comparison, when we analyzed sample replicates without OTCD, in the negative ionization mode, using NEDC (*N*-(1-naphthyl) ethylenediamine dihydrochloride) as a MALDI matrix, less than ten annotated molecular formulae were annotated with high confidence, and the list of annotations can be found on METASPACE (26). This comparison clearly demonstrates the necessity of alternative approaches for carbonyl detection from agar samples.

In the 4-APEBA-based OTCD analysis, several spatial patterns that depict changes in specific metabolite production between isolated and interacting microbes were observed (**Figure 1**). For example, we identified a group of molecules produced and excreted by *B. subtilis* only in its interaction with *Fusarium* sp. (*e.g.*, hexosamine, **Figure 1F**). Conversely, the production of an *N*-acetylated form of hexosamine (likely *GlcNAc*) was triggered in *Fusarium* sp., but not *B. subtilis*, with the interaction of these two species (**Figure 1C**). This finding is congruent with previous reports that *GlcNAc* acts as a signal inducer within fungi and also serves in interkingdom communication (27). In this case, it is possible that fungi utilize hexosamine produced by *B.subtilis* for GlcNAc biosynthesis but it is more likely that GlcNAc is released from fungi cell wall (chitin) during the interaction.(28) Contrasting spatial distributions were observed with citrate and homocitrate (**Figure 1G and 1D**, respectively), which are two chemically and metabolically related molecules. We observed homocitrate to be produced only in isolated *B. subtilis* colonies, while citrate, a central metabolite of the TCA cycle (29), was produced in both isolated and interacting *Fusarium* sp.. Based on citrate co-localization with *B. subtilis* cells only in the interaction zone, we hypothesize that citrate originating from *Fusarium* sp. might serve as a carbon source for adjacent *B. subtilis* colonies during the interaction. Our MSI data also found that other carbonyls have a variety of unique spatial patterns, including those: unchanged during interaction (*e.g.*, succinyl-glutamate, **Figure 1E**), present in distinct phenotypes of the *B. subtilis* biofilm (*e.g.*, acetyl-citrulline, **Figure 1H**), suppressed in both species when interacting (*e.g.*, aminobutanoate, **Figure 1I**), and activated in both species in their interactions (*e.g.*, acetamidopentanoate, **Figure 1J**). A list of all other metabolites annotated, together with their discriminant coefficients (AUC) between isolated and interacted species, can be found in **Supplementary Table S1.**

**Figure 1.**
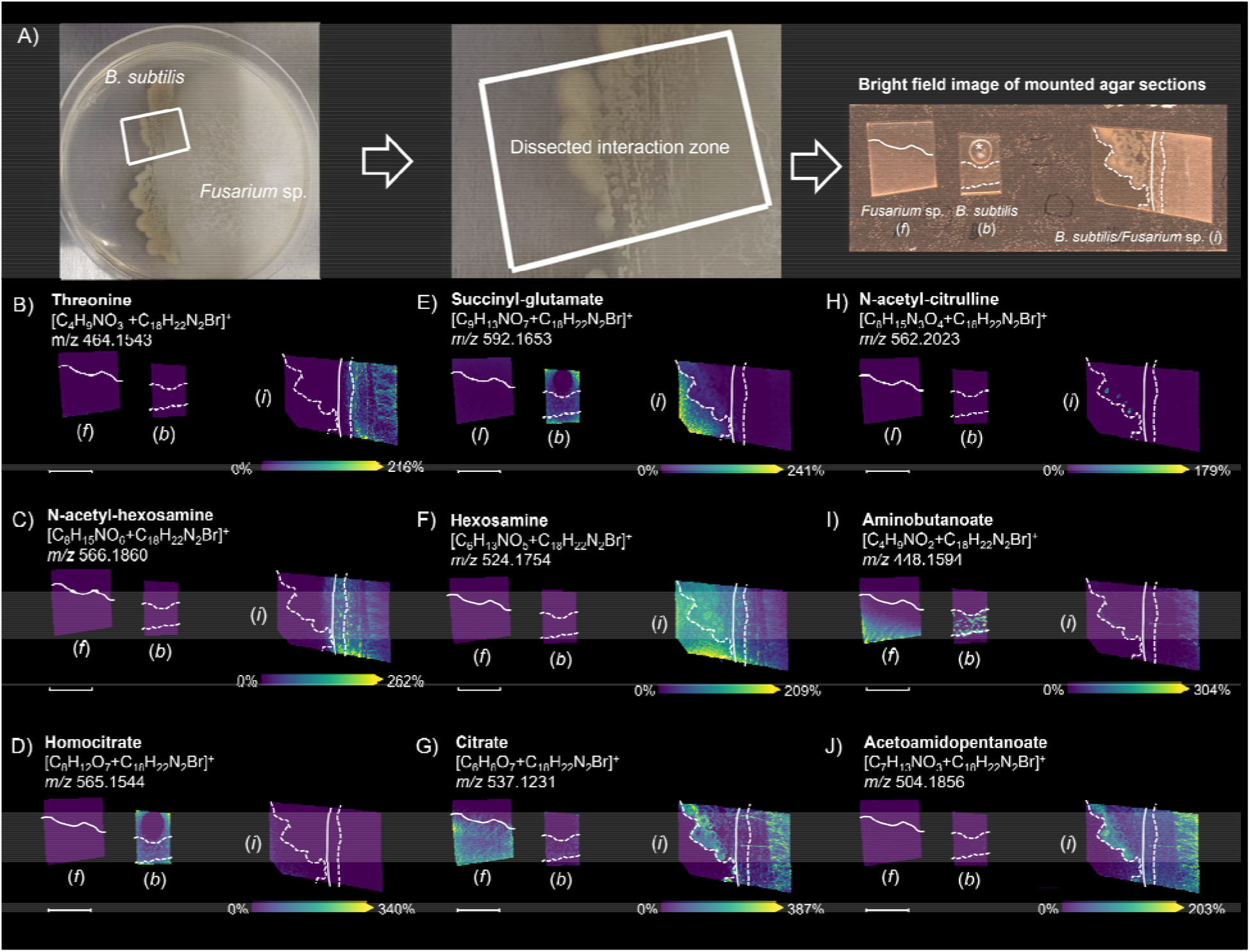
Characteristic patterns observed in the production and distribution of carbonyls from the interaction of *B. subtilis* and *Fusarium* sp. using the 4-APEBA-based OTCD approach. Each image is annotated with (*f*) showing the isolated *Fusarium* sp. Control, (*b*) showing the isolated *B. subtilis* control, and (*i*) showing the interaction zone of the co-culture of *B. subtilis* and *Fusarium* sp. **A)** Photograph of B. subtilis and Fusarium sp. in interactingon on the agar plate, a zoomed view into dissected interaction zone, and an optical image of isolated colonies and the interaction of B. subtilis and Fusarium sp. Colonies colonies mounted on the double-sided copper tape covered slide. MALDI-MSI ion images of carbonyls are highlighted that show: in **B)** and **C)** increased production in *Fusarium* sp. while in interaction with *B. subtilis*; **D)** suppressed excretion from *B. subtilis* while in interaction with *Fusarium* sp.; **E)** no change in abundance between isolated and colonies in interaction; **F)** increased excretion from *B. subtilis* during interaction with *Fusarium* sp.; **G)** increased production in *B. subtilis* and steady state in *Fusarium* sp. during interaction; **H)** hot spots in *B. subtilis* biofilm during interaction with *Fusarium* sp.; **I)** suppressed production in both *B. subtilis* and *Fusarium* sp. in interaction compared to isolated cultures; and **J)** increased production in both *B. subtilis* and *Fusarium* sp. in interaction compared to isolated culture. Solid and dashed white lines on the ion images indicate boundaries of *Fusarium sp*. and *B. subtilis* colonies, respectively. Scale bars are 7 mm and each ion images intensity is respectively scaled.**SMART** annotation:(44) **S** (step size, spot size, total scans) = 100 µm, 30 µm x 30 µm, 37,672 scans; **M** (molecular confidence) = MS1, 3 ppm; **A** (annotations) = 316 (METASPACE, KEGG (20% FDR), [M+C H N Br]^+^); **R** (resolving power) = 110,000 at *m/z* 400; **T** (time of acquisition) = 745 min.

Besides homocitrate (**Figure 1D**), citrate (**Figure 1G**), and several dozen other polycarboxylic acids (**Supplementary Table S1)**, we annotated numerous aliphatic monocarboxylic acids (*i.e.*, free fatty acids; FFA) as 4-APEBA derivatives (**Figure 2**). Their spatial profile points to the different roles of these molecules during the *B. subtilis* and *Fusarium* sp. interaction. For example, there is intense excretion of short-chain FFA (**Figure 2A-B**) from *B. subtilis* during interaction with *Fusarium* sp.. In contrast, medium-chain FFA (C14-C16, **Figure 2C-E**) are colocalized with *B. subtilis* cells further from the interaction zone. Strikingly, those with even number of carbons (C14 and C16) are also highly excreted in the surrounding agar in isolated bacteria and fungi cultures, whereas C15 are not observed in any of isolated cultures. It is known that *Bacillus* spp. can modify their FFA patterns to adapt to a wide range of environmental changes (30), and based on our results, it seems that the production of short-chain FFA is vital for surviving in a *Fusarium* sp. environment. On the other hand, *Fusarium* sp., in interaction with *B. subtilis* boosts the production of unsaturated long chain FFA (C18, octadecatrienoic acid and linoleate, **Figure 2F-G**), while saturated long chain FFA seem to be more significantly excreted from isolated colonies than within interaction (**Figure 2H**). This reinforced production of unsaturated long-chain FFA might be related to the antimicrobial defense mechanisms of *Fusarium* sp.. Namely, the antibacterial properties of FFA are used by many organisms, where the prime target of FFA action is the bacterial cell membrane, where FFA disrupts the electron transport chain and oxidative phosphorylation (31). Optical microscopy images of co-cultured *B. subtilis* NCIB 3610 and *Fusarium* sp. DS 682 demonstrate an antagonistic interaction as there is reduced fungal growth in presence of *B. subtilis* compared to the control (**Supplementary Figure S2**). *B. subtilis* is known to protect host plants by decreasing pathogenic fungal or bacterial growth through production of secondary metabolites (32), and as the majority of these compounds contain carbonyls, 4-APEBA-based OTCD enables the sensitive tracing of their redistribution and kinetics.

**Figure 2.**
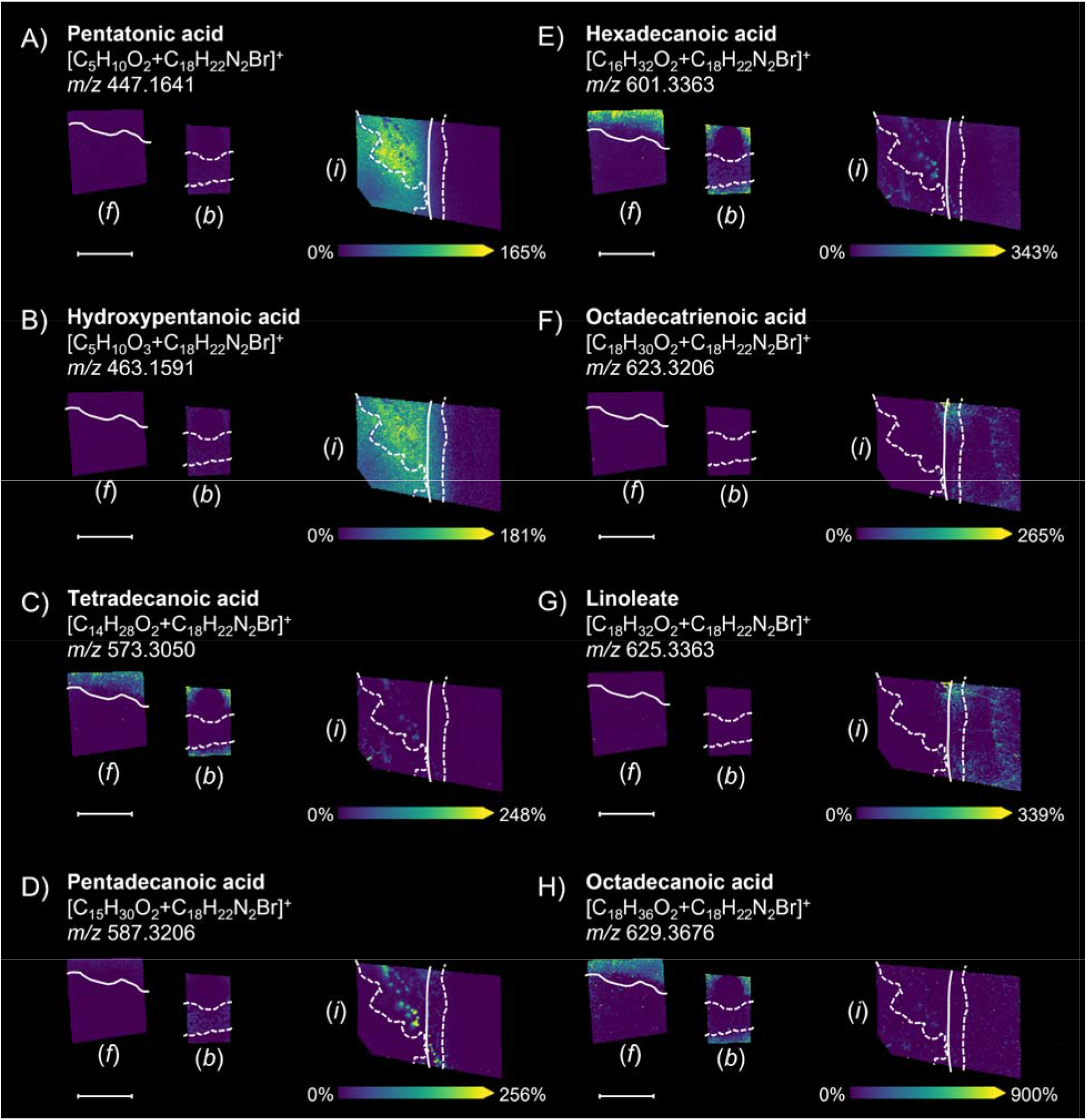
Shift in production and distribution of select aliphatic carboxylic acids during the *B. subtilis* and *Fusarium* sp. interaction, as well as isolated controls of each culture using the 4-APEBA-based OTCD approach. Each image is annotated with (*f*) showing the isolated *Fusarium* sp. Control, (*b*) showing the isolated *B. subtilis* control, and (*i*) showing the interaction zone of the co-culture of *B. subtilis* and *Fusarium* sp.. Solid and dashed white lines on the ion images indicate boundaries of *Fusarium sp*. and *B. subtilis* colonies, respectively. Scale bars are 7 mm and each ion images intensity is respectively scaled. **SMART** annotation:(44) **S** (step size, spot size, total scans) = 100 µm, 30 µm x 30 µm, 37,672 scans; **M** (molecular confidence) = MS1, 3ppm; **A** (annotations) = 316 (METASPACE, KEGG (20% FDR), [M+C H N Br]^+^); **R** (resolving power) = 110,000 at *m/z* 400; **T** (time of acquisition) = 745 min.

Besides comprehensive carbonyl coverage, the additional value of this approach is the ability to confidently resolve some isomeric and isobaric metabolites (**Figure 3**) that are undistinguishable in typical MALDI-MSI experiments, which only provides accurate mass measurements without gas phase separation by ion-mobility. **Figure 3A** shows a MALDI-MS ion image of a brominated derivative product ion at *m/z* 560.1754, illustrating that this metabolite is concentrated on the *B. subtilis* biofilm layer further from the interaction zone. Based on the accurate mass measurements, this ion, within a 3 ppm window, can be ascribed to the derivatized forms of either kinetin (C _10_H_9_N_5_) or succinyl proline (C _9_H_13_NO_5_) (**Figure 3C**). Both molecules are naturally present in soil (*e.g.*, as a plant hormone and common amino acid, respectively). Since kinetin does not possess a carbonyl group that can be derivatized with 4-APEBA, it suggests that this ion corresponds to succinyl proline, as succinyl proline contains multiple carbonyl groups (**Figure 3C**). Interestingly, upregulation of succinylation is a known phenomenon in *B. subtilis* response to carbon source changes, especially when citrate becomes the carbon source (33). This succinylation-citrate relationship orthogonally validates our previous hypothesis that *Fusarium* sp. produces citrate (**Figure 1G**), which is further metabolized by *B. subtilis*.

**Figure 3.**
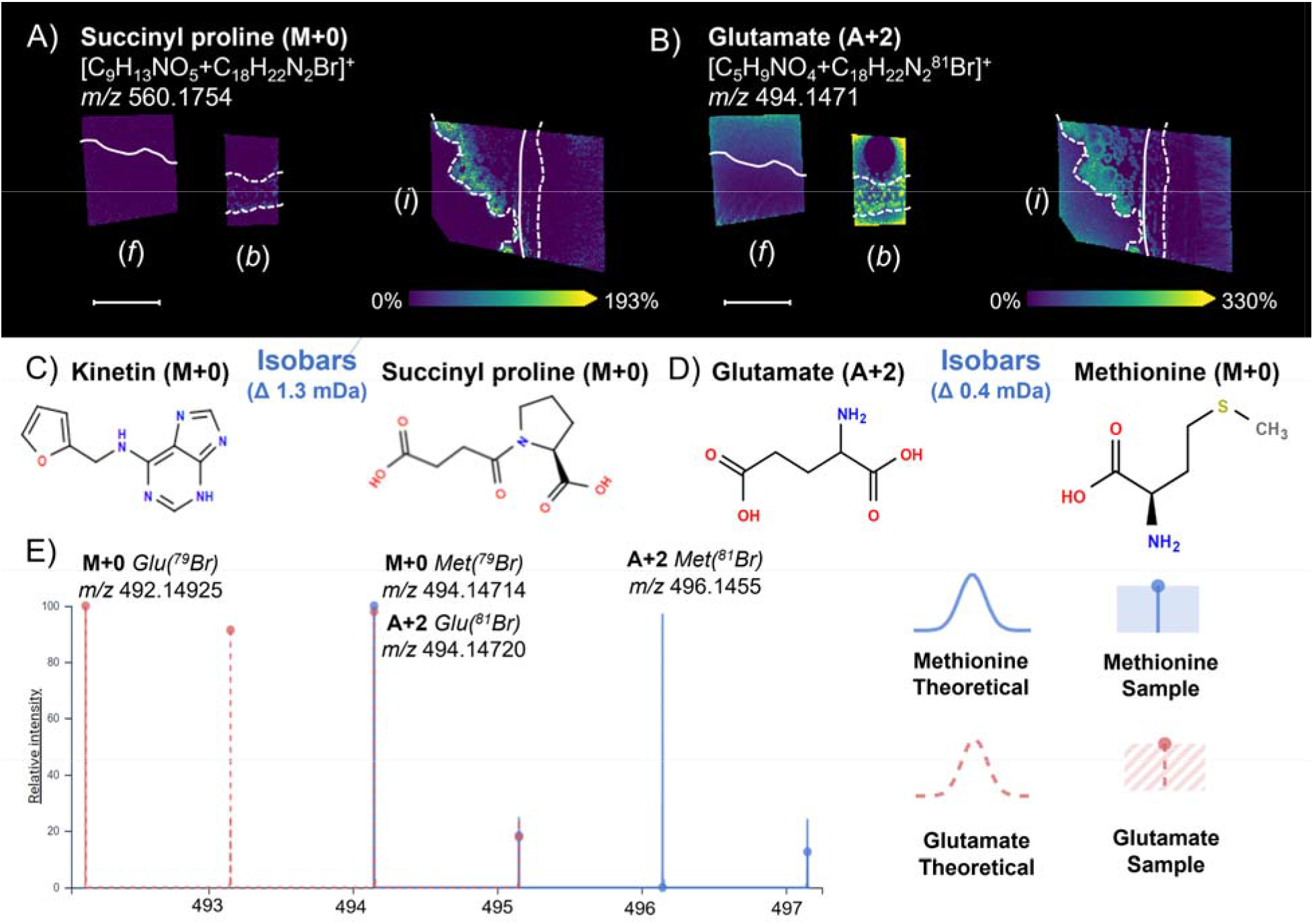
Resolving isobars using the 4-APEBA-based OTCD approach. Each image is annotated with (*f*) showing the isolated *Fusarium* sp. Control, (*b*) showing the isolated *B. subtilis* control, and (*i*) showing the interaction zone of the co-culture of *B. subtilis* and *Fusarium* sp. **A)** MALDI-MS ion images of *m/z* 560.1754 are shown with **C)** chemical structures of tentatively annotated derivatized isobaric metabolites monoisotopic kinetin (M+0, C_10_H_9_N_5_O+C_18_H_22_N_2_^79^Br) and monoisotopic succinyl proline (M+0, C_9_H_13_NO_5_+C_18_H_22_N_2_^79^Br), which differ by 1.3 mDa. Since kinetin does not contain a carbonyl that can derivatized, this confirms the annotation of succinyl proline. B) MALDI-MS ion images of *m/z* 490.1471 are shown with **D)** chemical structures of tentatively annotated derivatized isobaric metabolites monoisotopic methionine (M+0, C_5_H_11_NO_2_S+C_18_H_22_N_2_^79^Br) and the second isotopologue glutamate (A+2, C_5_H_9_NO_4_+ C_18_N_2_H_22_^81^Br), which differ by 0.4 mDa. **E)** Theoretical simulations and sample spectrum of isotopic distributions of both tentatively annotated isobars of methionine and glutamate from METASPACE, where within the sample spectrum the monoisotopic (M+0, ^79^Br) and the third (A+2, ^81^Br) isotopologue of methionine are not present at ratios representative of ^79^Br and ^81^Br isotopic distributions (blue trace vs blue dot). This confirms methionine as a false annotation of the third isotopologue (A+2, ^81^Br) of glutamate. Solid and dashed white lines on the ion images indicate boundaries of *Fusarium sp*. and *B. subtilis* colonies, respectively. Scale bars are 7 mm and each ion images intensity is respectively scaled. **SMART** annotation:(44) **S** (step size, spot size, total scans) = 100 µm, 30 µm x 30 µm, 37,672 scans; **M** (molecular confidence) = MS1, 3ppm; **A** (annotations) = 316 (METASPACE, KEGG (20% FDR), [M+C _18_H_22_N_2_Br]^+^); **R** (resolving power) = 110,000 at *m/z* 400; **T** (time of acquisition) = 745 min.

The second example of resolving isobars with 4-APEBA-based OTCD illustrates the importance of using bromine as a non-leaving moiety of the derivatization agent. Namely, bromine has two stable isotopes (^79^Br and ^81^Br) with similar relative abundances (51% and 49%, respectively), producing an easily recognizable isotopic pattern, where the monoisotopic (M+0; ^79^Br) and second isotopologue (A+2; ^81^Br) peaks have similar intensities. For instance, the ion at *m/z* 494.1471 (**Figure 3B**) can be wrongly annotated as methionine (M+0; ^79^Br derivative) but is actually an isotopologue of glutamate (A+2; ^81^Br derivative). If this were methionine (**Figure 3D**), an A+2 isotopologue at *m/z* 496.1455 would have a similar intensity as the putative monoisotopic peak at *m/z* 494.1471 (**Figure 3E**), which was not the case. Instead, the peak at *m/z* 492.1492 has the same intensity and exact spatial localization as *m/z* 494.1471, indicating glutamate was derivatized.

Lastly, the two-step 4-APEBA-based OTCD approach could be utilized to also differentiate some carbonyl isomers (**Figure 4**). This is because if 4-APEBA is used alone,(34) then it can only derivatize aldehydes and ketones, and it will not derivatize carboxylic acids (21). EDC must be used as an activator for 4-APEBA to derivatize carboxylic acids (**Supplementary Figure 3**).(35, 36) As such, we performed 4-APEBA-based OCTD with and without EDC on a replicate sample. Multiple overlapping and unique annotations were observed with side-by-side annotations outputs and ion images are visualized in METASPACE (37), and **Supplementary Table S2**. Overlapping annotations between both conditions (with and without the application of EDC) indicate the presence of a ketone or aldehyde group in the metabolite. Whereas in the case which the annotation was only present when EDC was applied prior to 4-APEBA, annotated molecules contain solely carboxylic acids, and aldehyde or ketone groups are absent from their structure and should not be considered. One example of an ambiguous annotation is the ion image at *m/z* 433.1476 (C _4_H_8_O_2_) (**Figure 4C-D**). This molecular formula can correspond to two natural products of bacterial metabolism: butanoic acid and acetoin (3-hydroxy-2-butanone) (**Figure 4G**). As a similar spatial pattern was observed with the addition of EDC (**Figure 4C**) and without the addition of EDC (**Figure 4D**), this ion image, which indicates intense excretion of the metabolite from *B. subtilis*, is likely is the ketone, acetoin, because the carboxylic acid, butanoic acid, cannot be derivatized without EDC. This annotation aligns well with the fact that acetoin is a primary catabolic product of *B. subtilis,* which bacteria reuse during the stationary phase when other carbon sources have been depleted (38). Moreover, the ion image at *m/z* 447.1641 with a similar spatial pattern was previously annotated as pentanoic acid (**Figure 2A**) and was not detected in the analysis without EDC (35), which orthogonally confirms the absence of butanoic acid in the analyzed sample. Other aliphatic carboxylic acids discussed throughout the manuscript (*i.e.,* FFA and some components of the TCA cycle) were also annotated only if EDC was used as an activator, while oxocarboxylic acids, due to the presence of ketone or aldehyde groups, were annotated without EDC as well (**Supplementary Table S2**). For example, the ion at *m/z* 479.1176 **(Figure 4E-F**) with the molecular formula of C _4_H_6_O_5_ is malate (two carboxyl groups, no ketone or aldehyde groups) rather than dehydro-threonate (carboxyl and ketone group), as this ion was not detected without EDC treatment (**Figure 4H**).

**Figure 4.**
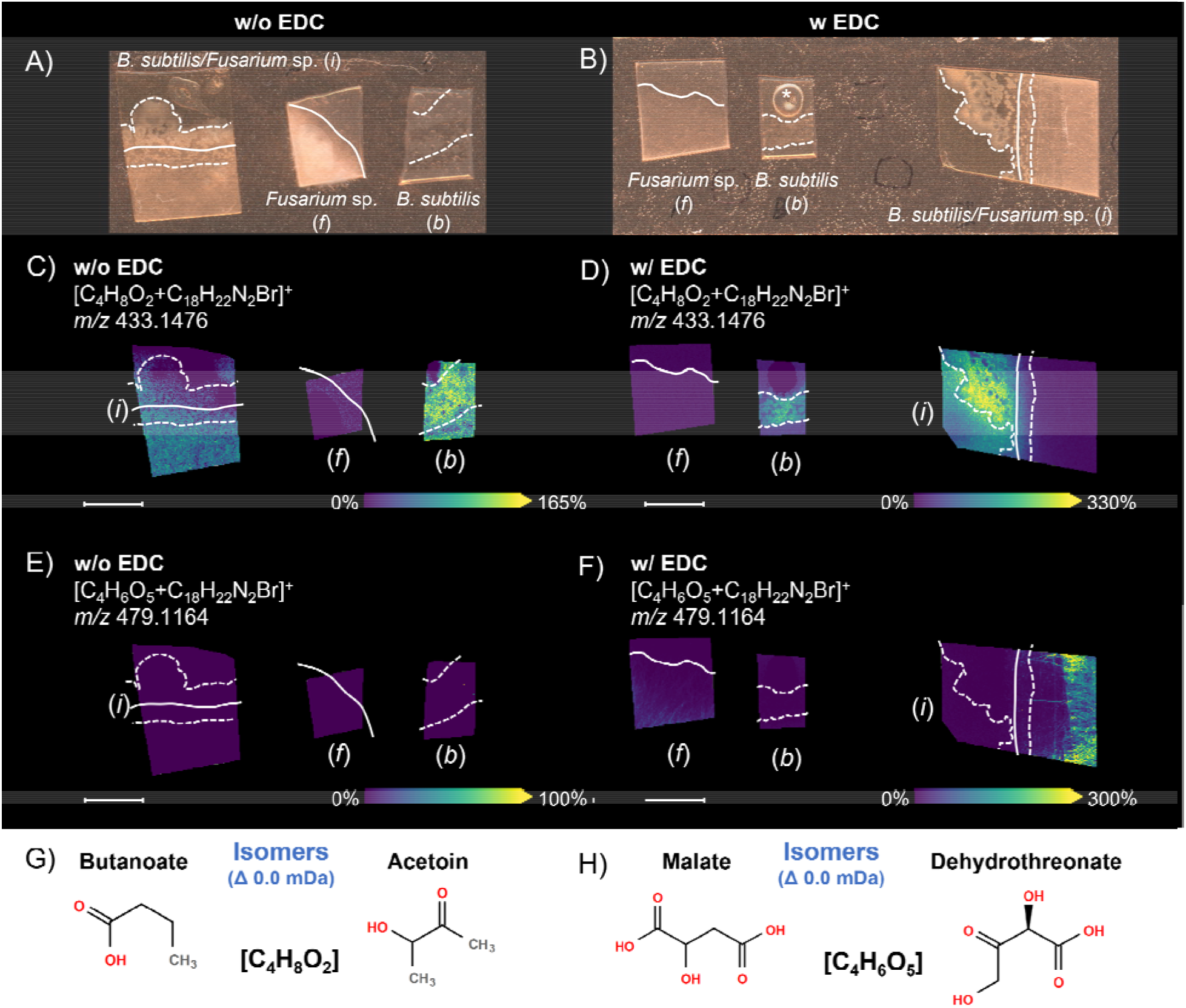
Resolving isomers using 4-APEBA-based OTCD with and without prior addition of EDC, which activates carboxylic acids prior to derivatization. Each image is annotated with (*f*) showing the isolated *Fusarium* sp. Control, (*b*) showing the isolated *B. subtilis* control, and (*i*) showing the interaction zone of the co-culture of *B. subtilis* and *Fusarium* sp.. Brightfield images from samples which underwent preparation **A)** without EDC and **B)** with EDC. MALDI-MSI ion images are shown for *m/z* 433.1476 **C)** without EDC and **D)** with EDC. Additional MALDI-MS ion images of *m/z* 479.1164 are shown **E)** without EDC and **F)** with EDC. Here, the ion at *m/* 479.1164 was not detected within **E)** without prior activation of carboxylic acids with EDC. **G)** The molecular structures of two tentatively annotated isomers, butanoic acid (carboxylic acid) and acetoin (ketone), are possible annotations for *m/z* 433.1476. **H)** The molecular structures of the other two tentatively annotated isomers, malate (solely carboxylic acid) and dehydrothreonate (oxoacid) are depicted for annotations at *m/z* 479.1164. Solid and dashed white lines on the ion images indicate boundaries of *Fusarium sp*. and *B. subtilis* colonies, respectively. Scale bars are 7 mm and each ion images intensity is respectively scaled. **SMART** annotation:(44) **S** (step size, spot size, total scans) = 100 µm, 30 µm x 30 µm, 37,672 scans; **M** (molecular confidence) = MS1, 3 ppm; **A** (annotations) = 316 (METASPACE, KEGG (20% FDR), [M+C_18_H_22_N_2_Br]^+^); **R** (resolving power) = 110,000 at *m/z* 400; **T** (time of acquisition) = 745 min.

This EDC-guided selectivity of 4-APEBA was additionally confirmed by analysis of several standards with different chemistries. Specifically, citric acid, which solely has carboxyl groups; glyoxalic acid, which has a carboxylic and aldehyde group; hydroxyacetone, which has only a ketone group (analog to acetoin); and pyruvic acid, which has both a carboxylic and ketone group (**Figure 5**). Our results show that citric acid, with no ketone or aldehyde groups, was not derivatized without the activation by EDC.(36) Due to the presence of a ketone in pyruvic acid, and an aldehyde group in glyoxalic acid these oxoacid metabolites were derivatized without EDC, although with ∼50% lower signal intensities than with EDC. This difference in intensity could be a consequence of matrix heterogeneity in dried droplet preparations, but might also be resultant from different derivatization efficiency between the two treatments. In fact, hydroxyacetone, the demethylated analog of acetoin, which we previously distinguished from its isobar butanoic acid (**Figure 4G**), shows clear isotopic patterns corresponding to its 4-APEBA derivatized product, and its signal is 2-fold higher without EDC than with EDC Therefore, we hypothesize that there is different kinetics and equilibrium in derivatization of ketoacids and ketones by 4-APEBA which can be exploited further for their differentiation. Importantly, none of the analytical standards show evidence of double or multiple derivatizations in any MALDI-FTICR mass spectra presented.

**Figure 5.**
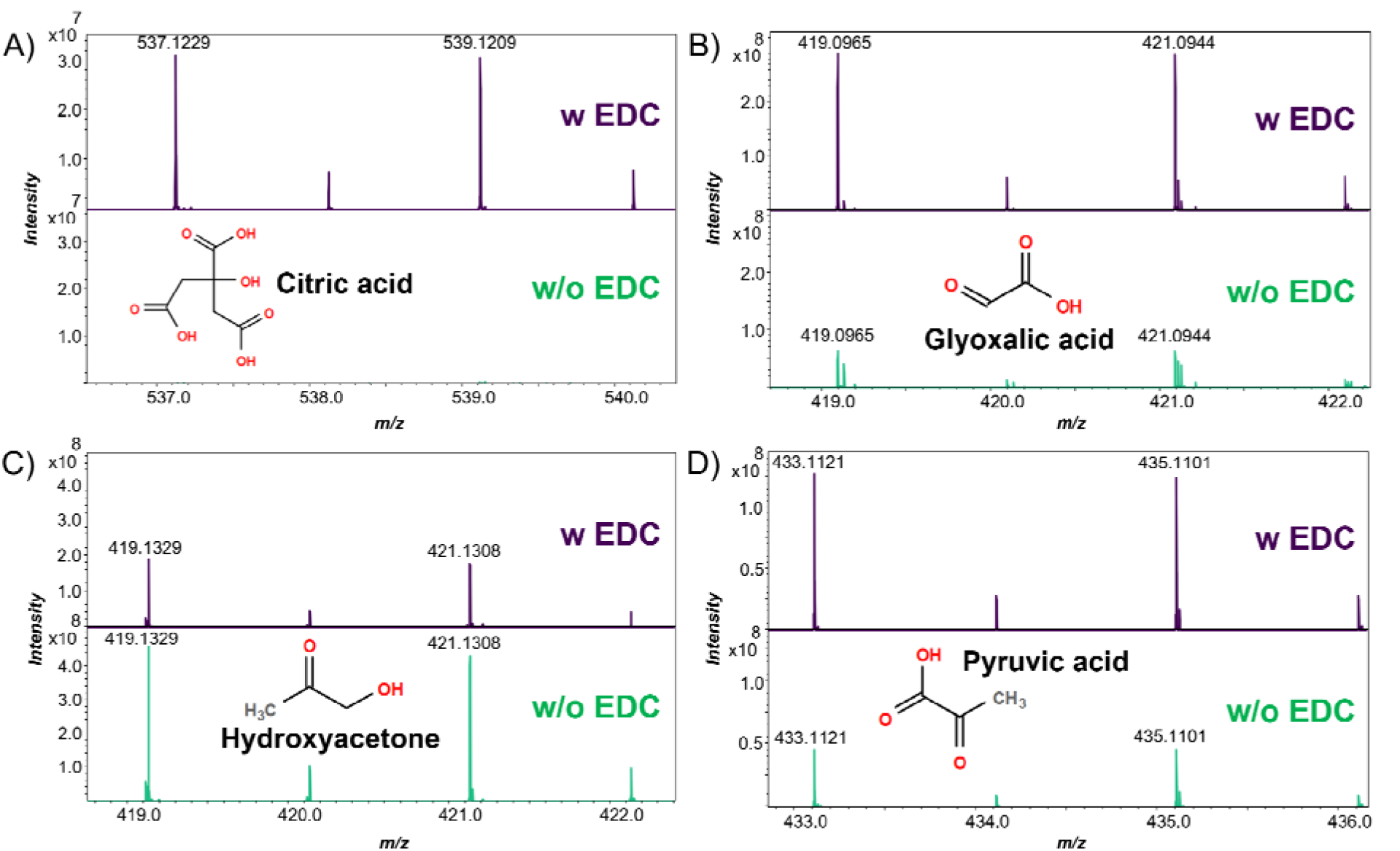
MALDI mass spectra of carbonyl standards with and without EDC activation prior to 4-APEBA derivatization. Mass spectra are shown for **A)** citric acid as a model carboxylic acid, **B)** glyoxalic acid as an oxocarboxylic acid with an aldehyde, **C)** hydroxyacetone as an oxoacid with a ketone, and **D)** pyruvic acid as an oxocarboxylic acid with a ketone. Peaks that correspond the monoisotopic (M+0, Br) and third isotopologue (A+2, Br) were annotated and shown in each spectrum and the structure of each metabolite is also presented.

## CONCLUSIONS

Mapping redistribution of acidic compounds in the microbial ecosystem using MALDI-MSI is a significant bioanalytical challenge. This challenge is further exacerbated for microbial cultures on agar, because many of these metabolites readily ionize in negative ionization mode which is not suitable for this type of sample. This work demonstrates a new approach for spatially profiling a diverse set of carbonyls, from small volatile aldehydes, ketones, and short-chain FFA to long-chain aliphatic carboxylic acids, including lactones and oxocarboxylic acids, directly from microbial cultures using 4-APEBA-based OTCD and MALDI-MSI. From our proof-of-concept experiments, we were able to map and reveal the distribution of carbonyls resulting from the interaction of *B. subtilis* and *Fusarium* sp. cultured on agar. These results illustrate the potential role of citrate, hexosamines, and FFA in this interaction. The additional advantage of the described OTCD by 4-APEBA, is that it can distinguish some isobaric and isomeric species, especially those with only a carboxyl group and only a ketone or aldehyde group, as well as those that do not have said functional groups at all. This benefit provides greater confidence in the biological interpretation of microbial MSI data. We also envision application of this workflow to map metabolic processes across a broad range of microbiology fields. For example, it can be used to spatially resolve the metabolome of digestive processes of the human gut, where microbiota produce large quantities of aliphatic acids and aldehydes through anaerobic fermentation of dietary fibers, which influences gut-brain communication and brain function (39). Another potential use is in resolving temporal-spatial chemistry of microbiological lignocellulose decay mechanisms where aldehydes, ketones, and carboxylic acids are both main degradation products and enzyme mediators for further lignin decomposition (40).

## MATERIAL AND METHODS

### Microbial growth and co-culture preparation

*B. subtilis* NCIB 3610 was cultured as described before (41). Briefly, the strain was cultured on MSgg, a *B. subtilis* biofilm-promoting medium (5 mM potassium phosphate [pH 7]; Fisher Scientific, Waltham, MA), 100 mM MOPS (morpholinepropanesulfonic acid [pH 7]; Sigma-Aldrich, St. Louis, MO), 2 mM MgCl_2_ (Fisher Scientific, Waltham, MA), 700 μM CaCl_2_ (Alfa Aesar, Haverhill, MA), 50 μM MnCl_2_ (Fisher Scientific, Waltham, MA), 50 μM FeCl_3_ (Sigma-Aldrich, St. Louis, MO), 11μM ZnCl_2_ (Sigma-Aldrich, St. Louis, MO), 2 μM thiamine hydrochloride (Fisher Scientific, Waltham, MA), 0.5 % glycerol (Fisher Scientific, Waltham, MA), and 0.5% glutamate (Sigma-Aldrich, St. Louis, MO). 1.5 % agar (BD) was added to the medium to prepare agar plates using MSgg media, and approximately 5 mL MSgg agar per plate were poured. MSgg agar plates were streaked with *B. subtilis* from frozen stocks and incubated at 28 °C for 15 hr.

*Fusarium* sp. DS 682 was cultured as described before (42). Fungi was maintained on potato dextrose agar (PDA) plates at 28 °C. For co-culture experiments a disposable 1.5 mm holepunch (Integra Miltex, York, PA) was used to collect the fungal biomass from a fungal culture PDA agar plate (1 agar plug with fungi) and was placed on the MSgg agar plate. Fungi was grown at 28°C for 5 days. After 5 days of fungal growth, *B. subtilis* was streaked close to the fungal biomass as described above. Agar plates were prepared for MSI analysis after 2 days of bacterial growth at 28°C.

Agar areas with isolated and interaction colonies were excised from MSgg agar Petri dishes, placed onto double-sided adhesive copper tape (3-6-1182; 3M United States) adhered to indium tin oxide-coated glass slides (Bruker Daltonics, Billerica, MA), and dried at RT overnight prior to derivatization and analysis (**Figure 1A**).

### **In-situ** chemical derivatization and MALDI matrix application

Agar samples were chemically derivatized by either sole application of synthesized 4-APEBA at 2 mg/mL or with a two-step approach: spraying an aqueous solution of EDC (1-ethyl-3-(3-dimethylaminopropyl)carbodiimide; Sigma-Aldrich, St. Louis, MO) at 6 mg/mL first with subsequent application of 4-APEBA at 2 mg/mL using an external syringe pump with the M5-Sprayer (HTX Technologies, Chapel Hill, NC). Spraying parameters were the same for both chemicals: 25 µL/min flow rate, a nozzle temperature of 37.5 °C, four cycles at 3 mm track spacing with a crisscross pattern, a 2 s drying period, 1,200 mm/min spray head velocity, 10 PSI of nitrogen gas, and a 40 mm nozzle height. DHB (2,5-dihydroxybenzoic acid; Sigma-Aldrich, St. Louis, MO) was prepared at a concentration of 40 mg/mL in 70% MeOH and was sprayed at 50 µL/min flow rate using the same M5-Sprayer. The nozzle temperature was set to 70 °C, with 12 cycles at 3 mm track spacing with a crisscross pattern. A 2 s drying period was added between cycles, a linear flow was set to 1,200 mm/min with 10 PSI of nitrogen gas and a 40 mm nozzle height. This resulted in matrix coverage of ∼667 µg/cm ^2^ for DHB. For negative ion mode experiments we used NEDC (N-(1-naphthyl)ethylenediamine dihydrochloride; Sigma-Aldrich, St. Louis, MO) which in our, and other laboratories,(43) yields more endogenous compounds than other commonly used matrices for negative mode analyses, such as 9-AA or 1,5-DAN. NEDC was prepared at a concentration of 7 mg/mL in 70% MeOH and was sprayed at 120 µL/min flow rate using the same M5-Sprayer. The nozzle temperature was set to 70 °C, with 8 cycles at 3 mm track spacing with a crisscross pattern. A 0 s drying period was added between cycles, a linear flow was set to 1,200 mm/min with 10 PSI of nitrogen gas and a 40 mm nozzle height. This resulted in matrix coverage of ∼187 µg/cm^2^ for NEDC.

### Derivatization of analysis of standards with and without EDC addition

Standards of citric acid (Sigma-Aldrich, St. Louis, MO), glyoxalic acid (Sigma-Aldrich, St. Louis, MO), hydroxyacetone (Sigma-Aldric, St. Louis, MO) and pyruvic acid (Sigma-Aldrich, St. Louis, MO) were prepared by dissolving each of them individually in miliQ water to final concentration of 0.01 mg/mL. For each EDC/4-APEBA reaction 10 µL of each standard was diluted in 400 µL of 6 mg/mL EDC and 400 µL of 2mg/mL 4-APEBA. For each 4-APEBA reaction without EDC, each standard was diluted in 400 µL milliQ water and 400 µL of 2 mg/mL 4-APEBA. Reactions were quenched after 2 hrs, and 1 µL of each reaction was spotted to a MALDI MTP 384 target plate (Bruker Daltonics, Billerica, MA) and mixed with 1 µL of DHB matrix (40 mg/mL in 70% MeOH).

### MALDI-MS analysis and data processing

All imaging and analyses of standards were performed on a Bruker Daltonics 12T solariX FTICR MS, equipped with a ParaCell, Apollo II dual ESI and MALDI source with a SmartBeam II frequency-tripled (355 nm) Nd: YAG laser (Bremen, Germany). Positive ion mode OTCD and negative ion mode NEDC acquisitions were acquired with broadband excitation from *m/z* 98.3 to 1,000, resulting in a detected transient of 0.5593 s — the observed mass resolution was ∼110k at *m/z* 400. FlexImaging (Bruker Daltonics, v.5.0) was used for the imaging experiments, and analyses were performed with a 100 µm step size. FlexImaging sequences were directly imported into SCiLS Lab (Bruker Daltonics, v.2023.a Premium 3D) using automatic MRMS settings. Ion images were directly processed from the profile datasets within SCiLS Lab, and automated annotation of the centroided dataset was completed within METASPACE with a chemical modifier corresponding to the mass shift expected from 4-APEBA derivatization (+C_18_H_22_N_2_Br, +345.0966 Da). KEGG-v1 was used as a metabolite database for annotations. All annotations were imported back to SCiLS as a new peak list, and discrimination analysis (ROC, receiver operation characteristic) between species in the interaction and corresponding isolated species were performed on that list. Area under the curve (AUC) of ROC analysis for each pair (isolated microbe vs microbe in interaction) was calculated.

### Data availability

All MALDI-MSI datasets and annotations are publicly available at METASPACE, a link for reviewers is here: https://metaspace2020.eu/api_auth/review?prj=b43e6772-d58c-11ed-afbb-53f7af38ceec&token=SVQu4QGZBzO6

## SUPPLEMENTAL MATERIAL FILE LIST

The following is included as supplementary material for this manuscript.

**Figure S1.** Supplemental file 1 contains this figure that shows the breakdown of assigned carbonyl derivative ions from KEGG-v1 output of METASPACE.

**Figure S2.** Supplemental file 1 contains this a figure that shows microscopy images of the growth of the co-cultured *B. subtilis* and *Fusarium* sp. on the MSgg agar medium.

**Figure S3.** Reaction scheme for derivatization of aldehydes/ketones and carboxylic acids with 4-APEBA.

**Table S1.** Supplemental file 2 is a complex workbook which shows the discriminate analyses completed for co-cultured *B. subtilis* and *Fusarium* sp. highlighting AUC within the isolated colonies and the interaction zone.

**Table S2.** Supplemental file 3 is a complex workbook which highlights all annotations from KEGG-v1 output of METASPACE for experiments with and without application of EDC before 4-APEBA.

## ACKNOWLEDGMENT

This research was performed on a project award doi.org/10.46936/intm.proj.2021.60091/60001441 (D.V.) from the Environmental Molecular Sciences Laboratory, a Department of Energy (DOE) Office of Science User Facility sponsored by the Biological and Environmental Research program under Contract No. DE-AC05-76RL01830. The funders had no role in study design, data collection and interpretation, or the decision to submit the work for publication.

## AUTHOR CONTRIBUTIONS

D.V. obtained funding and administered the project. D.V. and K.J.Z conceptualized the project. A.B. provided input into the microbiology and performed all microbiological work. A.B. created all cultures. D.V. performed derivatization and all MALDI-MSI experiments. D.V. and K.J.Z. performed data analysis and visualization. D.V. wrote the first draft of the manuscript with inputs from K.J.Z., A.B, and C.R.A. All authors edited the manuscript.

---

**Supplementary Figure S1.**
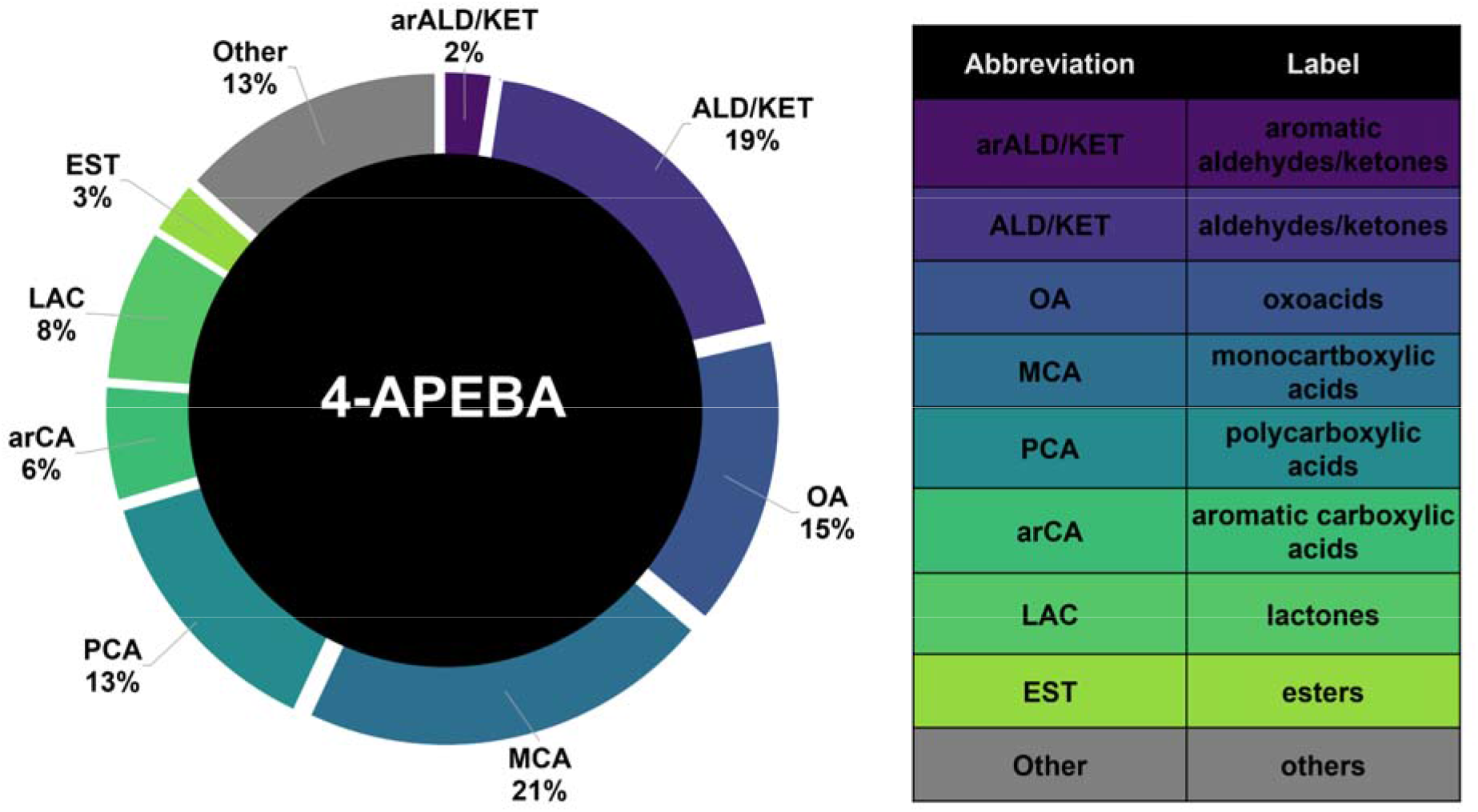
The breakdown of the carbonyl classes of the 316 assigned spectral peaks which underwent discriminate analysis within **Table S1** is shown. Here all annotation from KEGG-v1 output from METASPACE were bulked into representative carbonyl types, and then represented as a total percent of all bulked annotations of carbonyl type.

**Supplementary Figure S2.**
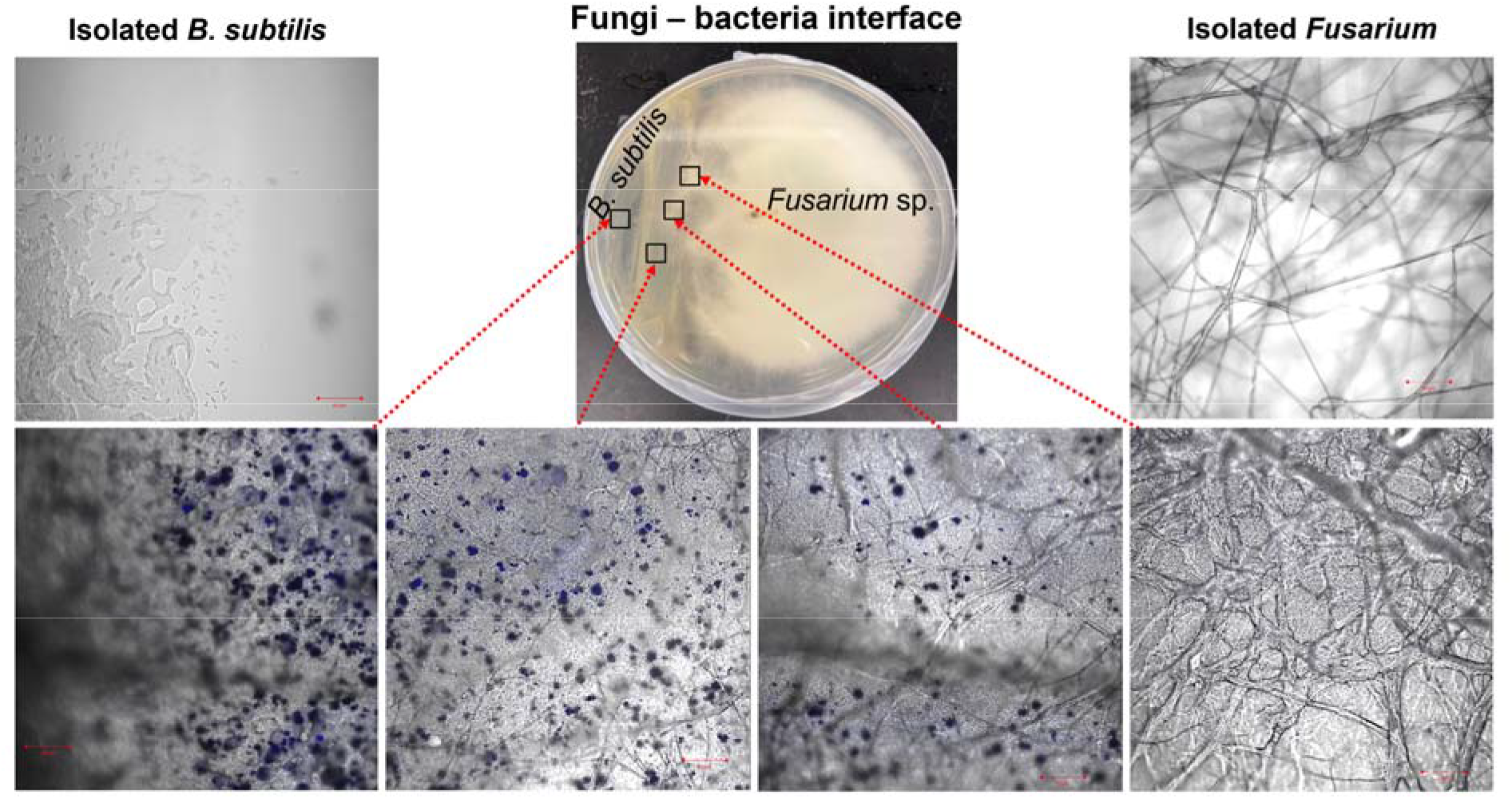
Optical microscopy images highlight microbial pigments produced a a result of fungi-bacteria interactions. Fungal hyphal architecture changes in presence of bacteria as shown in the inset zoomed views for the interaction zone (bottom panel) compared to isolated *B. subtilis* (top left) and *Fusarium* sp. (top right), all scale bars are 50 µm.

**Supplementary Figure S3.**
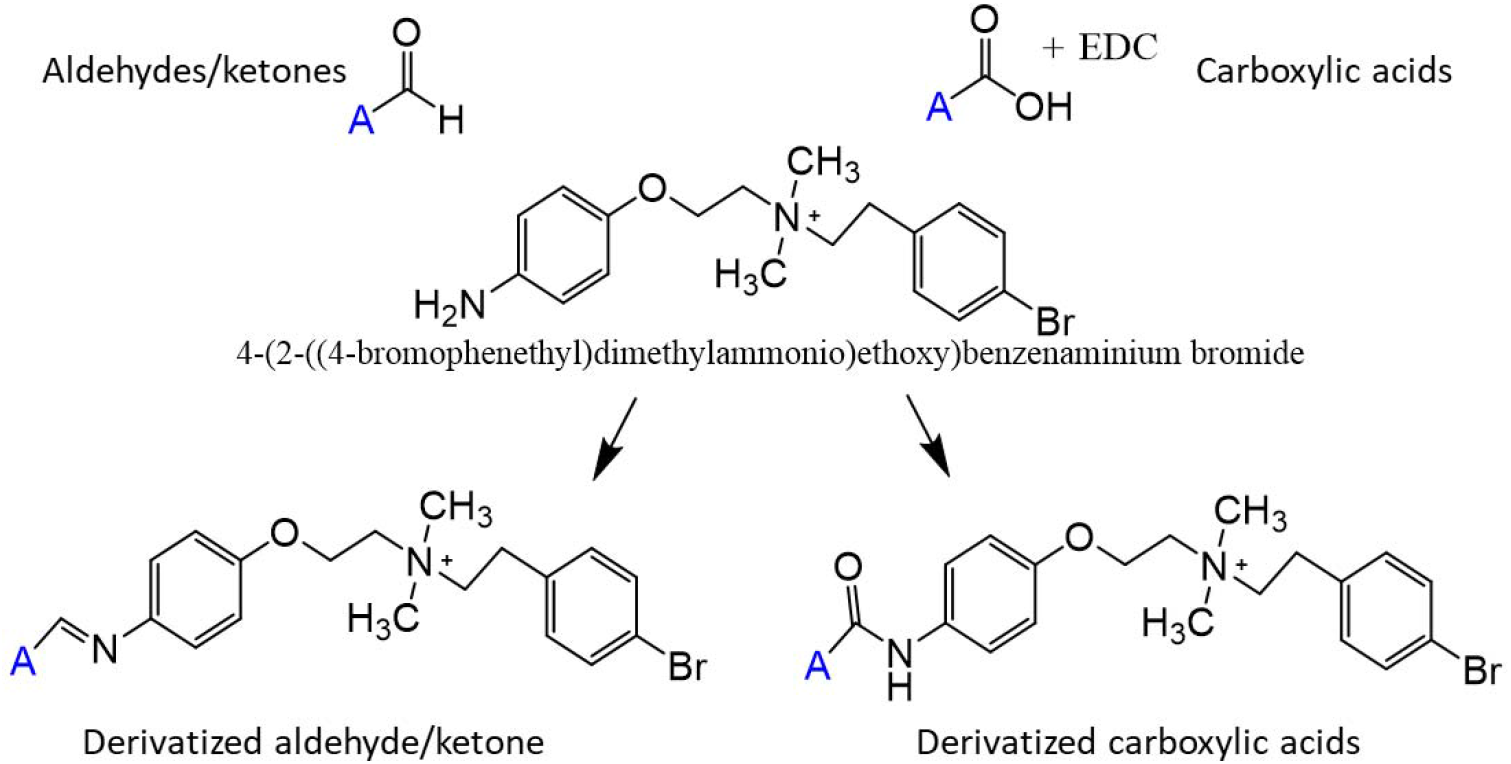
Reaction scheme for derivatization of aldehydes/ketones and carboxylic acids with 4-APEBA. Note that carboxylic acid needs to be activated with EDC before reaction with 4-APEBA.

## REFERENCES

1. Feucherolles M, Frache G. 2022. MALDI Mass Spectrometry Imaging: A Potential Game-Changer in a Modern Microbiology. Cells 11.

2. Dunham SJB, Ellis JF, Li B, Sweedler JV. 2017. Mass Spectrometry Imaging of Complex Microbial Communities. Accounts of Chemical Research 50:96–104.

3. Spraker JE, Luu GT, Sanchez LM. 2020. Imaging mass spectrometry for natural products discovery: a review of ionization methods. Natural Product Reports 37:150–162.

4. Yannarell SM, Velickovic D, Anderton CR, Shank EA. 2021. Direct Visualization of Chemical Cues and Cellular Phenotypes throughout Bacillus subtilis Biofilms. Msystems 6.

5. Stasulli NM, Shank EA. 2016. Profiling the metabolic signals involved in chemical communication between microbes using imaging mass spectrometry. Fems Microbiology Reviews 40:807–813.

6. Sgobba E, Daguerre Y, Giampa M. 2021. Unravel the Local Complexity of Biological Environments by MALDI Mass Spectrometry Imaging. International Journal of Molecular Sciences 22.

7. Watrous J, Roach P, Heath B, Alexandrov T, Laskin J, Dorrestein PC. 2013. Metabolic Profiling Directly from the Petri Dish Using Nanospray Desorption Electrospray Ionization Imaging Mass Spectrometry. Analytical Chemistry 85:10385–10391.

8. Li H, Balan P, Vertes A. 2016. Molecular Imaging of Growth, Metabolism, and Antibiotic Inhibition in Bacterial Colonies by Laser Ablation Electrospray Ionization Mass Spectrometry. Angewandte Chemie-International Edition 55:15035–15039.

9. Nagy G, Velickovic D, Chu RK, Carrell AA, Weston DJ, Ibrahim YM, Anderton CR, Smith RD. 2019. Towards resolving the spatial metabolome with unambiguous molecular annotations in complex biological systems by coupling mass spectrometry imaging with structures for lossless ion manipulations. Chemical Communications 55:306–309.

10. Musat N, Musat F, Weber PK, Pett-Ridge J. 2016. Tracking microbial interactions with NanoSIMS. Current Opinion in Biotechnology 41:114–121.

11. Boya CA, Fernandez-Marin H, Mejia LC, Spadafora C, Dorrestein PC, Gutierrez M. 2017. Imaging mass spectrometry and MS/MS molecular networking reveals chemical interactions among cuticular bacteria and pathogenic fungi associated with fungus-growing ants. Scientific Reports 7.

12. Pessotti RD, Hansen BL, Zacharia VM, Polyakov D, Traxler MF. 2019. High Spatial Resolution Imaging Mass Spectrometry Reveals Chemical Heterogeneity Across Bacterial Microcolonies. Analytical Chemistry 91:14818–14823.

13. Jens JN, Breiner DJ, Phelan VV. 2022. Spray-Based Application of Matrix to Agar-Based Microbial Samples for Reproducible Sample Adherence in MALDI MSI. Journal of the American Society for Mass Spectrometry 33:731–734.

14. Velickovic D, Chu RK, Carrell AA, Thomas M, Pasa-Tolic L, Weston DJ, Anderton CR. 2018. Multimodal MSI in Conjunction with Broad Coverage Spatially Resolved MS2 Increases Confidence in Both Molecular Identification and Localization. Analytical Chemistry 90:702–707.

15. Wu HM, Wu LK, Zhu Q, Wang JY, Qin XJ, Xu JH, Kong LF, Chen J, Lin S, Khan MU, Amjad H, Lin WX. 2017. The role of organic acids on microbial deterioration in the Radix pseudostellariae rhizosphere under continuous monoculture regimes. Scientific Reports 7.

16. Kunjapur AM, Prather KLJ. 2015. Microbial Engineering for Aldehyde Synthesis. Applied and Environmental Microbiology 81:1892–1901.

17. Ellis EM. 2002. Microbial aldo-keto reductases. Fems Microbiology Letters 216:123–131.

18. Parsons JB, Yao JW, Frank MW, Jackson P, Rock CO. 2012. Membrane Disruption by Antimicrobial Fatty Acids Releases Low-Molecular-Weight Proteins from Staphylococcus aureus. Journal of Bacteriology 194:5294–5304.

19. Tran VG, Zhao HM. 2022. Engineering robust microorganisms for organic acid production. Journal of Industrial Microbiology & Biotechnology 49.

20. Velickovic D, Lin VS, Rivas A, Anderton CR, Moran JJ. 2020. An approach for broad molecular imaging of the root-soil interface via indirect matrix-assisted laser desorption/ionization mass spectrometry. Soil Biology & Biochemistry 146.

21. Zemaitis KJ, Lin VS, Ahkami A, Winkler T, Anderton CR, Veličković D. 2023. [Preprint] Expanded Coverage of Phytocompounds by Mass Spectrometry Imaging Using On-Tissue Chemical Derivatization by 4-APEBA. ChemRxiv doi:https://doi.org/10.26434/chemrxiv-2023-v0xj4-v2.

22. Zhou QQ, Fulop A, Hopf C. 2021. Recent developments of novel matrices and on-tissue chemical derivatization reagents for MALDI-MSI. Analytical and Bioanalytical Chemistry 413:2599–2617.

23. Larson EA, Forsman TT, Stuart L, Alexandrov T, Lee YJ. 2022. Rapid and Automatic Annotation of Multiple On-Tissue Chemical Modifications in Mass Spectrometry Imaging with Metaspace. Analytical Chemistry 94:8983–8991.

24. Sejalon-Cipolla M, Bruyat P, Bregant S, Malgorn C, Devel L, Subra G, Cantel S. 2021. Targeting out of range biomolecules: Chemical labeling strategies for qualitative and quantitative MALDI MS-based detection. Trac-Trends in Analytical Chemistry 143.

25. Bhattacharjee A, Anderson LN, Alfaro T, Porras-Alfaro A, Jumpponen A, Hofmockel KS, Jansson JK, Anderton CR, Nelson WC. 2021. Draft Genome Sequence of Fusarium sp. Strain DS 682, a Novel Fungal Isolate from the Grass Rhizosphere. Microbiol Resour Announc 10.

26. Dataset provided by Dušan Veličković EMSL, Pacific Northwest National Laboratory. Agar_NEDC, on METASPACE. https://metaspace2020.eu/annotations?db_id=374&prj=b43e6772-d58c-11ed-afbb-53f7af38ceec&ds=2023-03-13_22h46m34s. Accessed 04/27/2023.

27. Konopka JB. 2012. N-acetylglucosamine (GlcNAc) functions in cell signaling. Scientifica (Cairo) 2012.

28. Min K, Naseem S, Konopka JB. 2019. N-Acetylglucosamine Regulates Morphogenesis and Virulence Pathways in Fungi. J Fungi (Basel) 6.

29. Jung T, Mack M. 2018. Interaction of enzymes of the tricarboxylic acid cycle in Bacillus subtilis and Escherichia coli: a comparative study. Fems Microbiology Letters 365.

30. Diomande SE, Nguyen-The C, Guinebretiere MH, Broussolle V, Brillard J. 2015. Role of fatty acids in Bacillus environmental adaptation. Frontiers in Microbiology 6.

31. Desbois AP, Smith VJ. 2010. Antibacterial free fatty acids: activities, mechanisms of action and biotechnological potential. Applied Microbiology and Biotechnology 85:1629–1642.

32. Andrić S, Meyer T, Ongena M. 2020. Bacillus Responses to Plant-Associated Fungal and Bacterial Communities. Frontiers in Microbiology 11.

33. Kosono S, Tamura M, Suzuki S, Kawamura Y, Yoshida A, Nishiyama M, Yoshida M. 2015. Changes in the Acetylome and Succinylome of Bacillus subtilis in Response to Carbon Source. Plos One 10.

34. Dataset provided by Dušan Veličković EMSL, Pacific Northwest National Laboratory. Agar_w/o_EDC_w_4-APEBA, on METASPACE. https://metaspace2020.eu/annotations?ds=2023-03-31_23h15m39s&viewId=sU.~Vswu. Accessed 04/27/2023.

35. Dataset provided by Dušan Veličković EMSL, Pacific Northwest National Laboratory. Agar_w_EDC_w_4-APEBA, on METASPACE. https://metaspace2020.eu/annotations?ds=2023-03-08_02h49m09s&viewId=tNBAag-A. Accessed 04/27/2023.

36. Nakajima N, Ikada Y. 1995. Mechanism of amide formation by carbodiimide for bioconjugation in aqueous media. Bioconjugate chemistry 6:123–130.

37. Dataset provided by Dušan Veličković EMSL, Pacific Northwest National Laboratory. Comparison of Agar_w/o_EDC_w_4-APEBA and Agar_w_EDC_w_4-APEBAon METASPACE. https://metaspace2020.eu/datasets/2023-03-31_23h15m39s/comparison?db_id=374&chem_mod=%2BN2H22C18Br&add=%5BM%5D%2B&viewId=pMCQpSSh&sort=datasetcount&row=4. Accessed 04/27/2023.

38. Ali NO, Bignon J, Rapoport G, Debarbouille M. 2001. Regulation of the acetoin catabolic pathway is controlled by sigma L in Bacillus subtilis. Journal of Bacteriology 183:2497–2504.

39. Silva YP, Bernardi A, Frozza RL. 2020. The Role of Short-Chain Fatty Acids From Gut Microbiota in Gut-Brain Communication. Frontiers in Endocrinology 11.

40. Velickovic D, Zhou MW, Schilling JS, Zhang JW. 2021. Using MALDI-FTICR-MS Imaging to Track Low-Molecular-Weight Aromatic Derivatives of Fungal Decayed Wood. Journal of Fungi 7.

41. Lukowski JK, Bhattacharjee A, Yannarell SM, Schwarz K, Shor LM, Shank EA, Anderton CR. 2021. Expanding Molecular Coverage in Mass Spectrometry Imaging of Microbial Systems Using Metal-Assisted Laser Desorption/Ionization. Microbiology Spectrum 9:e00520–21.

42. Bhattacharjee A, Qafoku O, Richardson Jocelyn A, Anderson Lindsey N, Schwarz K, Bramer Lisa M, Lomas Gerard X, Orton Daniel J, Zhu Z, Engelhard Mark H, Bowden Mark E, Nelson William C, Jumpponen A, Jansson Janet K, Hofmockel Kirsten S, Anderton Christopher R. 2022. A Mineral-Doped Micromodel Platform Demonstrates Fungal Bridging of Carbon Hot Spots and Hyphal Transport of Mineral-Derived Nutrients. mSystems 7:e00913–22.

43. Wang J, Qiu S, Chen S, Xiong C, Liu H, Wang J, Zhang N, Hou J, He Q, Nie Z. 2015. MALDI-TOF MS Imaging of Metabolites with a N-(1-Naphthyl) Ethylenediamine Dihydrochloride Matrix and Its Application to Colorectal Cancer Liver Metastasis. Analytical Chemistry 87:422–430.

44. Xi Y, Sohn AL, Joignant AN, Cologna SM, Prentice BM, Muddiman DC. 2023. SMART: A data reporting standard for mass spectrometry imaging. Journal of Mass Spectrometry 58:e4904.

