## Supplemental Information for "Mass spectrometry imaging of natural carbonyl products directly from agar-based microbial interactions using 4-APEBA derivatization"

*Running title: Direct imaging of carbonyls in microbial interactions*

Dušan Veličković,^a^* Kevin J. Zemaitis,^a^* Arunima Bhattacharjee,^a^ Christopher R. Anderton^a^

Environmental Molecular Sciences Laboratory, Pacific Northwest National Laboratory, Richland, WA 99354^a^

**Table of Contents:**

**Figure S1.** Figure that shows the breakdown of assigned carbonyl derivative ions from KEGG-v1 output of METASPACE.

**Figure S2.** Figure that shows microscopy images of the growth of the co-cultured *B. subtilis* and *Fusarium* sp. on the MSgg agar medium.

**Figure S3.** Reaction scheme for derivatization of aldehydes/ketones and carboxylic acids with 4-APEBA.

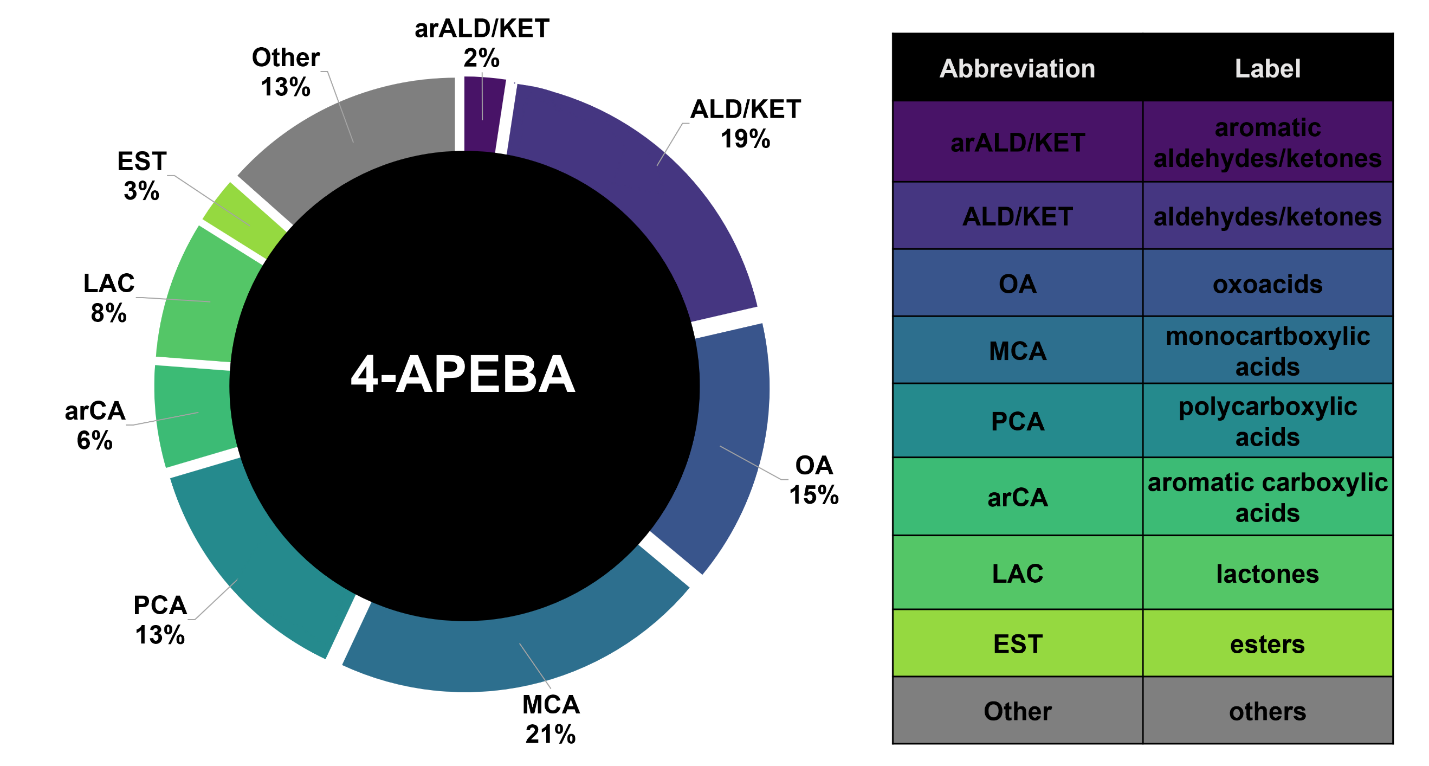

**Supplementary Figure S1.** The breakdown of the carbonyl classes of the 316 assigned spectral peaks which underwent discriminate analysis within **Table S1** is shown. Here all annotations from KEGG-v1 output from METASPACE were bulked into representative carbonyl types, and then represented as a total percent of all bulked annotations of carbonyl type.

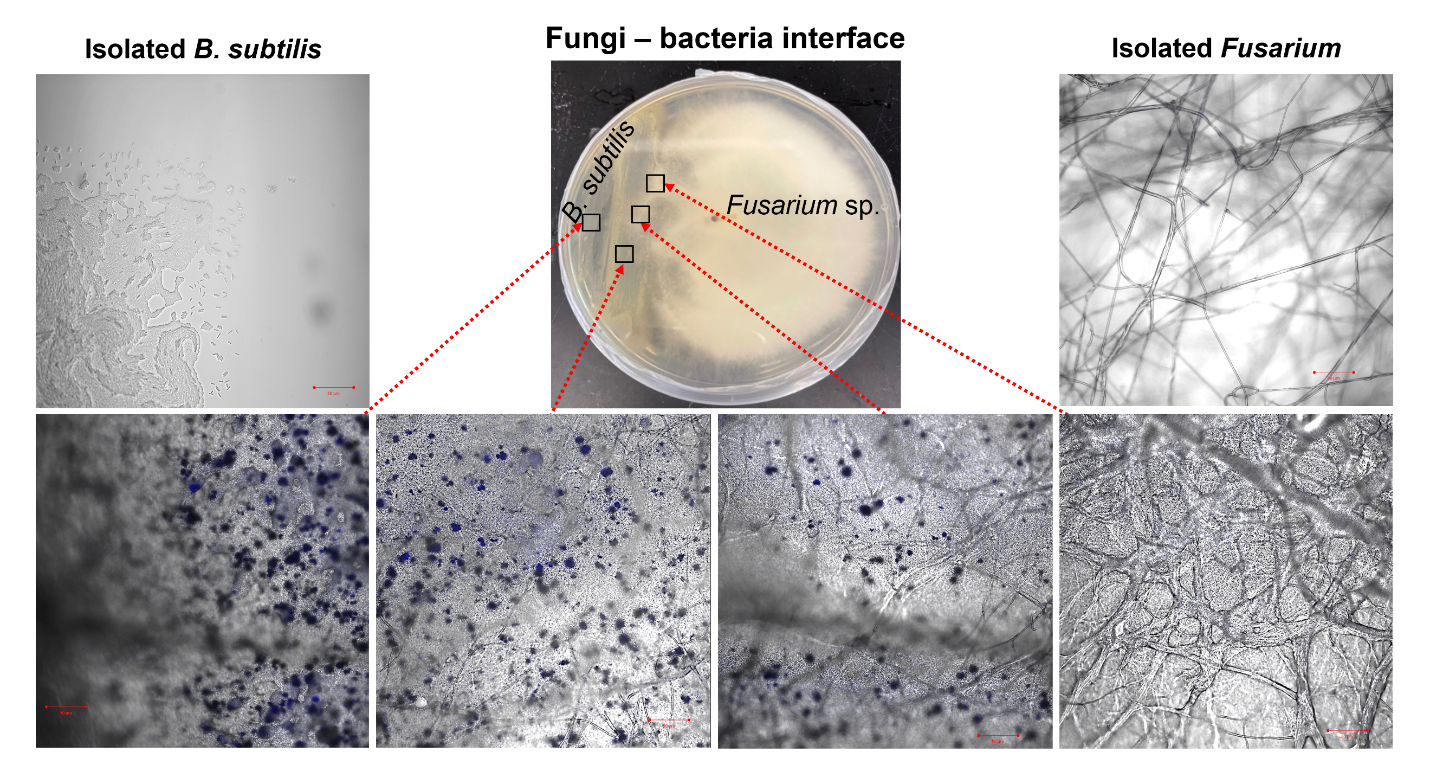

**Supplementary Figure S2.** Optical microscopy images highlight microbial pigments produced as a result of fungi-bacteria interactions. Fungal hyphal architecture changes in presence of bacteria as shown in the inset zoomed views for the interaction zone (bottom panel) compared to isolated *B. subtilis* (top left) and *Fusarium* sp. (top right), all scale bars are 50 µm.

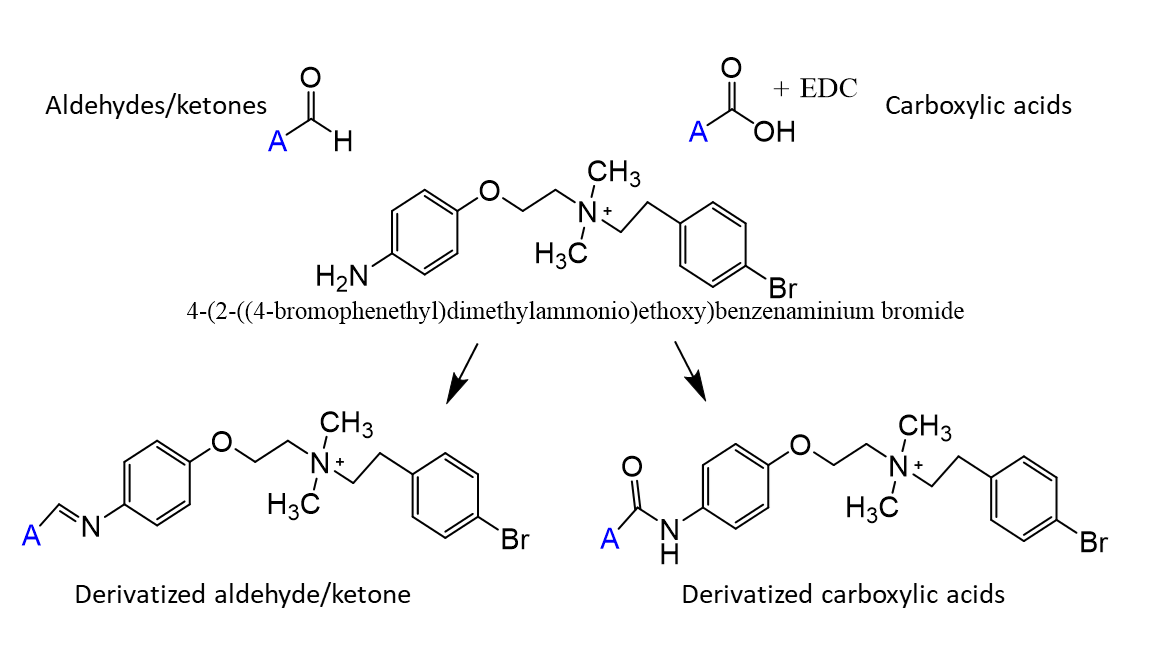

**Supplementary Figure S3.** Reaction scheme for derivatization of aldehydes/ketones and carboxylic acids with 4-APEBA. Note that carboxylic acid needs to be activated with EDC before reaction with 4-APEBA.
